# Widefield *in vivo* imaging system with two fluorescence and two reflectance channels, a single sCMOS detector, and shielded illumination

**DOI:** 10.1101/2023.11.07.566086

**Authors:** Patrick R. Doran, Natalie Fomin-Thunemann, Rockwell P. Tang, Dora Balog, Bernhard Zimmerman, Kıvılcım Kılıç, Emily A. Martin, Sreekanth Kura, Harrison P. Fisher, Grace Chabbott, Joel Herbert, Bradley C. Rauscher, John X. Jiang, Sava Sakadzic, David A. Boas, Anna Devor, Ichun Anderson Chen, Martin Thunemann

**Author notes:** **Corresponding authors:** Anna Devor and Martin Thunemann, 610 Commonwealth Ave, Boston, MA 02215, USA.

## Abstract

**Significance:** Widefield microscopy of the entire dorsal part of mouse cerebral cortex enables large-scale (“mesoscopic”) imaging of different aspects of neuronal activity with spectrally compatible fluorescent indicators as well as hemodynamics via oxy- and deoxyhemoglobin absorption. Versatile and cost-effective imaging systems are needed for large-scale, color-multiplexed imaging of multiple fluorescent and intrinsic contrasts.

**Aim:** Develop a system for mesoscopic imaging of two fluorescent and two reflectance channels.

**Approach:** Excitation of red and green fluorescence is achieved through epi-illumination. Hemoglobin absorption imaging is achieved using 525- and 625-nm LEDs positioned around the objective lens. An aluminum hemisphere placed between objective and cranial window provides diffuse illumination of the brain. Signals are recorded sequentially by a single sCMOS detector.

**Results:** We demonstrate performance of our imaging system by recording large-scale spontaneous and stimulus-evoked neuronal, cholinergic, and hemodynamic activity in awake head-fixed mice with a curved “crystal skull” window expressing the red calcium indicator jRGECO1a and the green acetylcholine sensor GRAB_ACh3.0_. Shielding of illumination light through the aluminum hemisphere enables concurrent recording of pupil diameter changes.

**Conclusions:** Our widefield microscope design with single camera can be used to acquire multiple aspects of brain physiology and is compatible with behavioral readouts of pupil diameter.

## Introduction

With the rapidly expanding list of fluorescent probes for neuronal activity and the development of large-scale measurements (Abdelfattah et al. 2022), widefield microscopy has become the measurement modality of choice for mesoscale studies of neuronal circuits and underlying behavior when cellular resolution is not required (Bouchard et al. 2009, Mohajerani et al. 2013, Ma et al. 2016b, Vanni et al. 2017, Wright et al. 2017, Couto et al. 2021). Probes available in distinct spectral variants (Zhao et al. 2011) can be spectrally multiplexed for simultaneous imaging of different aspects of neuronal activity (Lin and Schnitzer 2016). Using multiplexed probes for in vivo widefield imaging of head-fixed mice with “crystal skull” cranial windows (Kim et al. 2016) or transcranial imaging, optionally combined with thinned-skull preparations (Silasi et al. 2016), provides a view of the entire dorsal part of the cerebral cortex including the primary sensory, motor and higher association areas (Cramer et al. 2021, Higley and Cardin 2022).

In the healthy brain, increases in neuronal activity are often accompanied by increases in blood flow, volume, and oxygenation in the active region. The increase in blood flow and volume occurs, in large part, due to dilation of cerebral arterioles responding to vasoactive messengers released by active neurons (Uhlirova et al. 2016b, Echagarruga et al. 2020). This hemodynamic response increases oxygen supply beyond the actual demand leading to a net increase in blood oxygenation. Both oxy- and deoxyhemoglobin (HbO and HbR, respectively) strongly absorb light in the blue-green spectrum. Therefore, the hemodynamic response results in time-variant modulation of fluorescence signals within that spectral range. The methods developed to correct fluorescent signals for this hemodynamic artifact fall into two categories. For the first correction method, enhanced green fluorescent protein (EGFP)-based indicators, including GCaMP, iGluSnFR, and GRAB indicators, are excited off-peak at their isosbestic point in the 390-420 nm range. When excited at these wavelengths, fluorescence emission is independent from the analyte concentration (i.e., [Ca^2+^], [Glutamate], etc.), but affected by hemodynamic changes (Dana et al. 2019, Jing et al. 2020). This signal can then be regressed out of the analyte-dependent signal to remove hemodynamic artifacts. Correction methods measuring changes in hemoglobin absorption use either a single reflectance measurement at 520-530 nm or estimate changes in HbO and HbR concentrations (Δ□HbO] and Δ□HbR]) from simultaneous acquisition of light absorption from at least two wavelengths (Ma et al. 2016a). Furthermore, quantitative estimates of Δ[HbO] and Δ[HbR] are important measurables on their own; they aid in understanding cerebrovascular physiology and interpretation of non-invasive imaging signals generated through, for example, functional magnetic resonance imaging (fMRI) (Uhlirova et al. 2016a).

Current stationary widefield imaging setups typically use light-emitting diodes (LEDs) with sequential illumination, one or two low-noise, high-sensitivity scientific CMOS cameras, and low (smaller than 2x) magnification optics with a large field of view (Cardin et al. 2020). Experimental demands led to various adaptions of this basic blueprint. In systems dedicated for widefield calcium imaging with EGFP-based indicators, blue LEDs (450-480 nm) are used for fluorescence excitation, with light delivered through the objective lens or through side-illumination in some cases. Hemodynamic correction of green fluorescence signals either uses off-peak (390-420 nm) fluorescence excitation (Mohajerani et al. 2013, Lerner et al. 2015, Allen et al. 2017, Musall et al. 2019, Jacobs et al. 2020, Lake et al. 2020, MacDowell and Buschman 2020, Couto et al. 2021, Peters et al. 2021, Zatka-Haas et al. 2021) or is based on reflectance measurements at 520-530 nm (Wekselblatt et al. 2016, Vanni et al. 2017, Wright et al. 2017, Gilad et al. 2018, Mitra et al. 2018). Several groups incorporated a second fluorescence channel with excitation at 560-590 nm to enable two-color fluorescence imaging of red fluorescent calcium indicators like jRCaMP1b or jRGECO1a in combination with either EGFP-derived indicators such as GCaMP6 (Gribizis et al. 2019), GRAB_ACh_ (Lohani et al. 2022), or flavoprotein autofluorescence (Wang et al. 2022, Raut et al. 2023). Widefield imaging systems dedicated to the investigation of neurovascular interactions typically combine imaging of green (Ma et al. 2016b, Matsui et al. 2016, Wright et al. 2017, Mitra et al. 2018, Valley et al. 2020, Sunil et al. 2023), red fluorescent indicators (Park et al. 2020, Raut et al. 2023, Shahsavarani et al. 2023), or green and red fluorescent indicators (Wang et al. 2022) with reflectance imaging for Δ□HbO] and Δ□HbR] estimation at two or three wavelengths, typically in the range of 525-565 nm and 590-625 nm. Light for reflectance imaging is delivered directly from one or more multi-color light engines (Wright et al. 2017, Mitra et al. 2018, Shahsavarani et al. 2023), through a single light guide (Park et al. 2020), through an illumination ring (Harrison et al. 2009, White et al. 2011, Jacobs et al. 2020), or through several optical fibers connected to a single light source (Valley et al. 2020) to illuminate the sample surface at an angle. Some researchers report the use of linear polarizers in front of the light sources to minimize specular reflection from the glass window (White et al. 2011, Wright et al. 2017, Mitra et al. 2018). Experiments involving behavioral tasks or visual stimulation, different solutions to block illumination light from reaching the animal’s eyes have been reported, for example through implanted light shielding (Makino et al. 2017, Valley et al. 2020), custom light-shielding between objective and headplate (Couto et al. 2021, Nietz et al. 2023), or imaging stage design (Padawer-Curry et al. 2023).

Here, we describe a versatile widefield imaging system that enables two-color fluorescence imaging in addition to concurrent acquisition of two reflectance channels for quantitative estimation of Δ□HbO] and Δ□HbR] with a single sCMOS detector. Specular reflections from curved glass cranial windows that interfere with reflectance imaging are minimized using diffuse illumination with 525- and 625-nm light for reflectance imaging by combining a custom-built ring illuminator with an aluminum hemisphere between objective and cranial window. In addition, the aluminum hemisphere shields illumination light to prevent it from reaching the animal’s eyes and thereby avoiding unwanted visual stimulation of the animal. We provide optical design blueprints, a wiring diagram, a timing chart, data acquisition software, and an inventory of all parts. We demonstrate the performance of our new widefield microscope in awake mice expressing the mApple-based calcium indicator jRGECO1a (Dana et al. 2016) and the EGFP-based acetylcholine (ACh) probe GRAB_ACh3.0_ (Jing et al. 2020).

## Results

We designed and manufactured a widefield imaging system with a field of view of 10×10 mm2 covering the dorsal cortex of a mouse. We use a single sCMOS detector for sequential recording of two fluorescent indicators (here: EGFP-derived GRAB_ACh3.0_ and mApple-derived jRGECO1a) as well as hemoglobin absorption at 525 and 625 nm at an effective frame rate of 10 Hz or better without changes in the configuration of the emission path. **Figure 1** and **Supplementary Figure 1** provide an overview of the system layout and illustrate the spectral properties of the light sources and filter elements together with HbO and HbR absorption spectra as well as jRGECO1a and GRAB_ACh3.0_ excitation and emission spectra. For fluorescence excitation, we use epi-illumination through the objective lens. A 470-nm LED with a maximal power of 3.7 W provides light for GRAB_ACh3.0_ excitation, which is filtered through a 466/40 nm excitation bandpass filter, resulting in a maximal power of 403 mW at the sample surface. For jRGECO1a excitation, we use light from a 565-nm LED with a maximal power of 8.6 W which is filtered through a 560/14 nm excitation bandpass filter reaching a maximal power of 75 mW at the sample surface. A secondary peak in the 565-nm LED spectrum at ∼425 nm is removed by an additional 450-nm longpass filter in the excitation path (**Figure 1A**). The collimated and filtered light is coupled into the infinity space using a multiband dichroic mirror (488/561 nm, ‘main dichroic’). This, in combination with a multi-band ‘detection filter’ (523/610 nm), allows GRAB_ACh3.0_ and jRGECO1a fluorescence at 500-540 nm and 580-640 nm, respectively, to reach the sCMOS detector while blocking excitation light at 440-480 nm and 550-570 nm from reaching the camera (**Figure 1D, E, G**). Typical powers at the sample surface during image acquisition were 90-177 mW and 68-75 mW for 470 nm and 565 nm at illumination times of 3-6 ms, respectively. We observed no or minimal bleaching over continuous acquisition periods of up to 30 minutes. The wavelengths for hemoglobin absorption imaging are within the respective emission bands of the green (525 nm) and red (625 nm) fluorophore (**Figure 1F**). The LEDs used for hemoglobin absorption imaging are placed in a custom-designed reflectance illumination ring (RIR) located between the objective lens and the sample surface (**Figure 1B** and **Supplementary Figure 1D**). The RIR contains six 525-nm LEDs and six 625-nm LEDs positioned at equal distances with maximal powers of 170 mW and 371 mW per LED, respectively. To enable a simple procedure for estimation of Δ[HbR] and Δ[HbO], we placed 10-nm bandpass filters in front of each LED. This leads to a maximal power of 39.5 mW (525 nm) and 51 mW (625 nm) at the sample surface. During image acquisition, typical powers at the sample surface were 10.5 mW and 4.4 mW for the 525-nm and 625-nm LEDs at 3-ms illumination times.. We placed a hollow aluminum hemisphere between the RIR and the sample surface internally reflecting the light of the 525-nm and 625-nm LEDs. For imaging, the opening of the hemisphere is brought close to the sample surface achieving diffuse illumination and efficiently eliminating most reflections from the curved glass window covering the brain surface.

**Figure 1.**
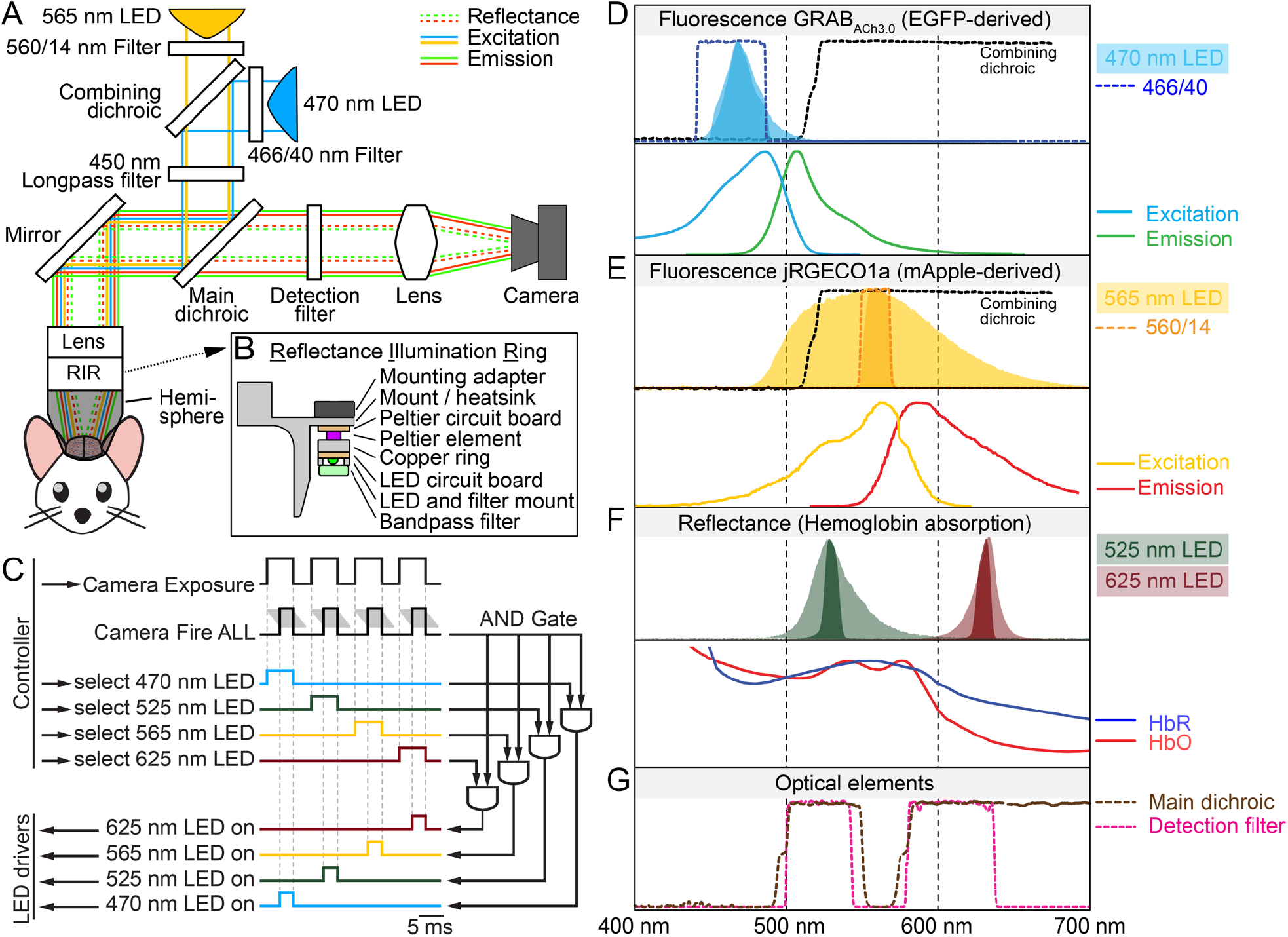
Schematics of the wide field imaging system and fluorescence and reflectance spectra. **A**. A single sCMOS camera is used to measure emission of green and red fluorophores and two hemoglobin absorption wavelengths. A 565-nm LED excites mApple-based jRGECO1a and a 470-nm LED excites EGFP-based GRAB_ACh3.0_; both LEDs are filtered through excitation filters. The main dichroic mirror reflects excitation light towards a 45°-mirror and then a 0.63× objective lens to focus the excitation light onto the sample. Below the objective lens is the reflectance illumination ring (RIR) further detailed in panel B. An aluminum hemisphere between the RIR and the sample surface internally reflects the 525- and 625-nm LEDs for diffuse illumination prevents specular reflections and illumination light from reaching the animal’s eyes. Fluorescence emission and reflectance pass the main dichroic and the detection filter before being focused by a tube lens to the camera. **B**. Cross-section of the reflectance illumination ring (RIR) setup. Each of the six 525- and 625-nm LEDs are arranged on a ring-shaped board at equal distances. Peltier elements actively cool the LEDs to prevent temperature changes resulting in unstable illumination intensities. 10-nm wide excitation filters are mounted in front of each LED. **C**. Timing diagram to trigger LEDs and camera acquisition: MATLAB controls the DAQ system via digital triggers controlling camera and LED drivers. A different LED illuminates the sample during each frame for near-simultaneous acquisition. The effective frame rate is determined by the length of the four-frame cycle (typically 100 ms). Grey lines next to the camera Fire ALL trigger represent exposure of pixel rows in rolling shutter mode. The Fire ALL trigger is sent when exposure overlaps for all rows. Each LED driver (bottom left) is controlled by the output of a separate AND gate. The Fire ALL trigger is the common input to every AND gate while the individual LED selection triggers are the second input for each AND gate. **D**. Top, light-blue and dark-blue shaded areas represent the 470-nm LED spectrum before and after passing the excitation filter (BP466/40, blue dotted line). Bottom, excitation (solid blue line) and emission (solid green line) spectrum of EGFP-derived GRAB_ACh3.0_ (Jing et al. 2020). **E**. Top, light-orange and dark-orange areas represent the 565-nm LED spectrum before and after passing the excitation filter (BP560/14, orange dotted line). Bottom, excitation (solid yellow line) and emission (solid red line) spectrum of mApple-derived jRGECO1a (Dana et al. 2016). **F**. Top, LED spectra are confined by excitation filters (525 nm in light and dark green; 625 nm in light and dark red before and after passing through the respective excitation filters). Bottom, absorption spectra of oxyhemoglobin (HbO, red line) and deoxyhemoglobin (HbR, blue line) from Prahl (1999). **G**. Spectra of the main dichroic (brown) and detection filter (pink). Light must pass through both filters to reach the camera.

With the chosen sCMOS camera working in rolling shutter mode, we ensure that the LEDs emit light only when all rows of the sCMOS chip are simultaneously exposed. Therefore, we control the LED drivers with two digital trigger signals combined through a logical AND gate integrated circuit (IC) (**Figure 1C**). The first input to the AND gate IC is generated by the camera and is in ‘high’-state when all rows are being exposed; the second input is generated by the microscope control computer and selects the respective LED wavelength to be switched on (**Figure 1C**). Only when both signals are in ‘high’ state, the LED power supply receives a trigger signal to switch the respective LED on. Timing delays of the AND gate IC and the LED driver are <9 ns and <100 µs, respectively, enabling virtually delay-free LED power switches for typical exposure times of 2-8 ms per individual LED.

The imaging setup is controlled through a DAQ interface programmed via MATLAB. The main acquisition script generates digital trigger signals controlling the imaging setup based on exposure time, which can be different for every wavelength, acquisition rate, and total acquisition time. Additional triggers are generated to synchronize or control external devices such as stimulus apparatuses, behavioral cameras, electrophysiology recording equipment, and behavioral control devices. The currents of the individual LEDs are set in MATLAB and sent via a USB-based remote-control interface to the LED drivers. The sCMOS camera is controlled through the manufacturer’s software where parameters such as field of view and on-camera pixel binning are set. This software also drives image acquisition; however, the camera is set to respond to exposure triggers generated by the DAQ interface, with the (variable) trigger length determining the exposure time. During acquisition, the DAQ interface records feedback signals from camera, microscope control, and external devices allowing synchronization of data streams during off-line post-processing and analysis.

While our single-camera approach reduces complexity and cost, we use multiband filters to image two fluorophores present in the sample at the same time. The lack of spectral separation on the emission side can potentially create crosstalk when fluorophores are excited off-resonance. Note that both fluorophores undergo dynamic changes in fluorescence intensity, so the degree of crosstalk will vary in time and space. Published spectra show that GRAB_ACh3.0_ (or EGFP) is essentially not excited at 565 nm, while jRGECO1a (or mApple) fluorescence is excited at 470 nm. To quantify the amount of crosstalk under well-controlled conditions, we collected images of purified EGFP and mApple proteins with various combinations of illumination power and exposure time for both the 470-nm and 565-nm LEDs. In line with the published spectra, we observed that the 565-nm LED practically does not excite purified EGFP while mApple shows fluorescence when excited with the 470 nm LED (**Supplementary Figure 2A** and **B**). Based on this *in vitro* calibration with EGFP and mApple and the typical acquisition parameters we used for *in vivo* imaging in GRAB_ACh3.0_- and jRGECO1a-expressing animals, we determined that the level of crosstalk of GRAB_ACh3.0_ to fluorescence measured upon 565-nm excitation is <1%, while fluorescence from jRGECO1a contributes about 3-9% to the fluorescence intensity measured upon 470-nm excitation (**Supplementary Figure 2**; see **Appendix B** for a detailed discussion of the crosstalk analysis). To verify that the degree of crosstalk is similar in recordings from awake mice expressing GRAB_ACh3.0_ and jRGECO1a, we applied a simplified correction approach that does not incorporate dynamic changes of hemoglobin absorption on fluorescence excitation and emission (details in **Appendix B**). After applying the correction, we used cross-correlation analysis to verify that the degree of crosstalk is within the same range as established under in-vitro conditions (**Supplementary Figure 3**). Here, it should be noted that high correlation between GRAB_ACh3.0_ and jRGECO1a signals is expected primarily due to their underlying physiological covariance (Lohani et al. 2022) and is only influenced to a small degree by optical crosstalk, as shown in the analysis. However, while the in-vitro calibration provides us with the parameters necessary to perform crosstalk correction of in-vivo data, we decided not to further use crosstalk correction for the recorded in vivo data as it likely will add additional noise without sufficient benefit – at least under the experimental conditions present in this study (see **Appendix B** for further discussion).

In addition, it has been reported for mApple-based calcium indicators like jRGECO1a that illumination with blue light leads to photo-switching, which causes a reversible, calcium-independent increase in fluorescence intensity (Dana et al. 2016). In a control experiment with an awake jRGECO1a-expressing mouse, we did not observe jRGECO1a photo-switching under 470-nm illumination conditions typical for two-color fluorescence imaging in GRAB_ACh3.0_- and jRGECO1a-expressing mice. However, when the 470-nm illumination intensity was increased by an order of magnitude compared to typical imaging conditions, we observed photo-switching of jRGECO1a (**Supplementary Figure 4**).

To demonstrate the capabilities of our new imaging system design under in-vivo conditions, we recorded spontaneous and stimulus-induced changes in neuronal calcium, ACh, and hemodynamic signals in awake head-fixed mice expressing jRGECO1a and GRAB_ACh3.0_ (**Figure 2**). Mice were implanted with a curved glass cranial window exposing both hemispheres of the dorsal cortex while leaving the bone in-between the two hemispheres above the superior sagittal sinus intact (Kilic et al. 2020) (**Figure 2A**). Images were registered to the Allen Mouse Common Coordinate Framework (Wang et al. 2020) to generate time courses from pre-defined regions of the dorsal cortex. Measurements of spontaneous activity show a high correlation between neuronal calcium and ACh time courses (**Figure 2B** shows right secondary motor cortex), which is consistent with published results (Lohani et al. 2022). We then recorded the response to a sensory stimulus consisting of a 2-s, 3-Hz air puff sequence (20 repetitions, 25 s inter-stimulus interval) delivered to the right whisker pad (**Figure 2C-E**). Ratio maps show neuronal activation and functional hyperemia in the contralateral barrel cortex (green ROI in calcium map, **Figure 2C**). The stimulus-averaged time course calculated from the contralateral barrel cortex shows that the imaging system operates at sufficient signal to noise ratio to record the neuronal calcium response to the six individual air puffs comprising one stimulus train (**Figure 2D**). Despite high trial-to-trial variability (**Figure 2E**), which is consistent with published widefield imaging studies in awake mice (Musall et al. 2019), responses to individual air puffs are still discernible in individual trial recordings. The time course of the hemodynamic response shows a broad peak and post-stimulus undershoot. The hemodynamic correction we apply to the green fluorophore compensates for darkening that occurs when hemoglobin levels increase. **Supplementary Figure 5** compares the trial averaged ACh time courses with and without correction for hemodynamic artifacts.

**Figure 2:**
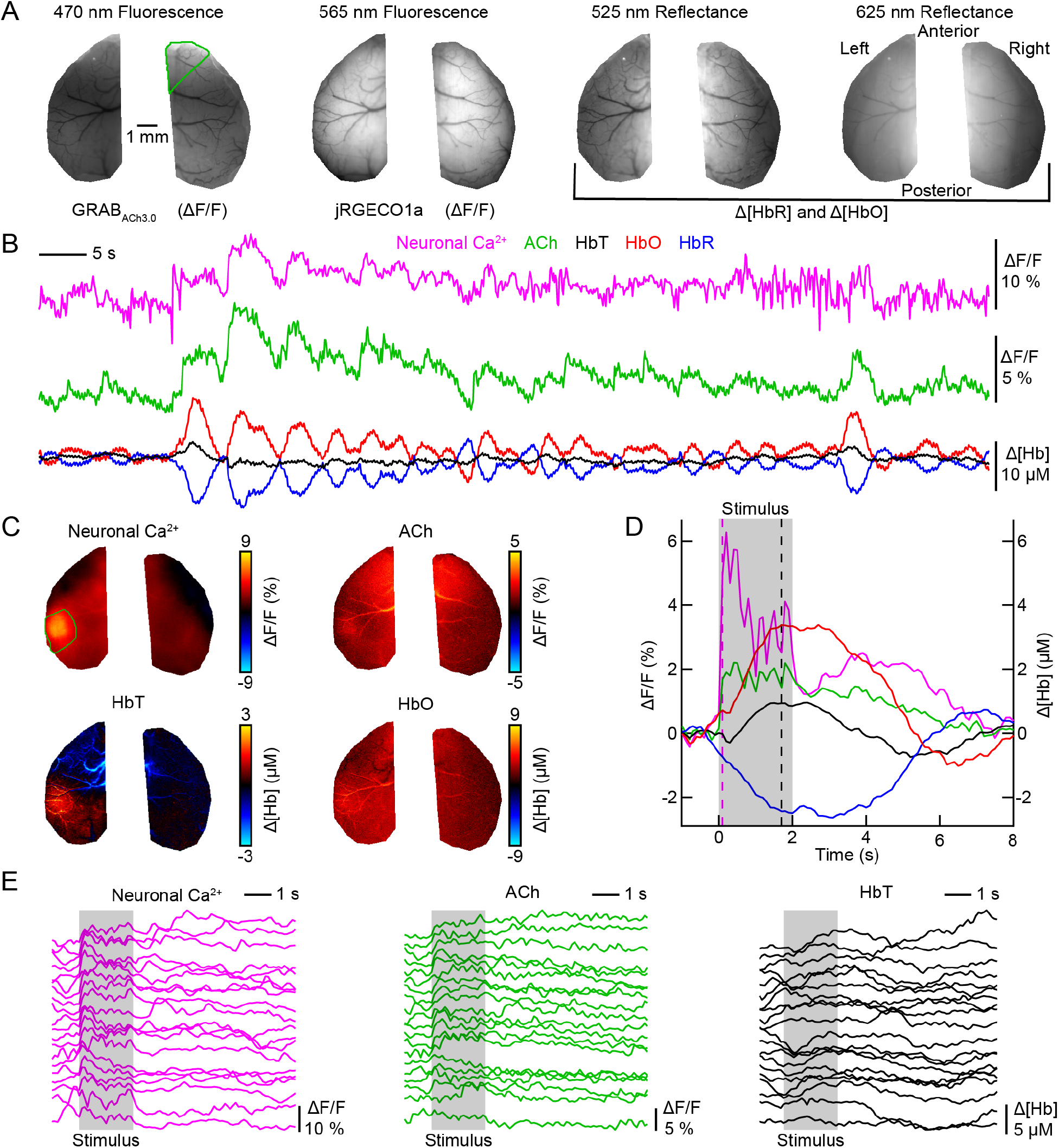
Spontaneous and stimulus-evoked activity in the somatosensory cortex. **A**. Average of the first ten images collected during a ten-minute acquisition for each illumination wavelength. The two hemispheres were manually masked around the cranial window. A green polygon was drawn around the right secondary motor cortex, which was determined by registration to the Allen Atlas of the mouse brain. **B**. Spontaneous activity in the right secondary motor cortex (green polygon in panel A) for calcium, acetylcholine (ACh), and oxygenated, deoxygenated, and total hemoglobin (HbO, HbR, HbT). **C**. Ratio maps averaged across 20 trials showing the response to a 2-s, 3-Hz train of air puffs to the right whisker pad. Maps show data from 1.7 s after stimulus onset except for the calcium map, which shows data from 0.1 s after stimulus onset. **D**. Average time course of the stimulus-evoked response in the contralateral barrel cortex (green region in panel C). The pink dashed line indicates when the calcium ratio map in panel C is shown whereas the black dashed line indicates when ACh, HbT and HbO ratio maps in panel C are shown. The grey-shaded area indicates the duration of the stimulus. **E**. Response of the contralateral barrel cortex to individual stimulus trains. The trials are sorted according to the magnitude of the calcium response.

With a unit price of >$20,000, the sCMOS camera is the most expensive component of the imaging system. To show how the system performance is affected by camera choice, we repeated the experiment shown in **Figure 2** with a CMOS camera costing aound $750 (**Supplementary Figure 6**). Time courses of signals averaged from spatially connected pixels in a region of interest show stimulus-evoked and spontaneous changes in neuronal calcium, ACh, and hemodynamic signals (**Supplementary Figure 6B, D, E**). On the other hand, we observed spatially structured noise across the field of view (compare **Figure 2C** with **Supplementary Figure 6C**) that could impact pixel-based analysis methods.

The aluminum hemisphere in-between the cranial window and the imaging system reflects 525- and 625-nm light and provides diffuse illumination minimizing specular reflections at the curved glass window that interfere with reflectance imaging. Inhomogeneities of signal intensity across the sample surface are likely occur due to the curvature of the ‘crystal skull’ glass window (**Figure 2A**). While enabling diffuse illumination, the aluminum hemisphere also acts as a shield preventing the strobing illumination light from reaching the mouse’s eyes. For example, four-color imaging with an effective acquisition rate of 10 Hz would produce an undesired visual stimulus at 40 Hz. To illustrate the shielding effect, we recorded changes of the animal’s pupil diameter during mesoscopic imaging in presence or absence of the aluminum hemisphere (**Figure 3**). For this experiment, a behavioral camera recorded the pupil diameter continuously while widefield illumination across the four wavelengths at 40 Hz was switched on and off at 60-s intervals; we recorded 20 off-to-on and on-to-off transitions per test. The images shown in **Figure 3A** show the pupil before and after turning the illumination on; **Supplementary Video 1** shows a recording of the off-to-on transition with and without hemisphere. **Figure 3B** shows representative 2-min time series covering the off-to-on and on-to-off transitions in presence or absence of the hemisphere. Without the hemisphere, pupil size measurements are largely dominated by the state of widefield illumination. In the presence of the hemisphere, pupil size changes are largely independent from widefield illumination (**Figure 3C**). Then, we compared the relative change in pupil size from 1.5-0.5 s before versus 2-3 s after start of widefield illumination averaged over 20 transitions and observe a significantly larger drop in pupil size without the hemisphere (p<0.001; t-test; **Figure 3D**). Pupil tracking is often used as an important variable to define behavioral states (Vinck et al. 2015, Reimer et al. 2016) and it is therefore desirable to combine pupil recordings and mesoscale imaging (Shahsavarani et al. 2023). In our system, the hemisphere eliminates interference of widefield illumination on behavioral recordings and enables high-quality measurements of pupil dynamics during mesoscale imaging.

**Figure 3:**
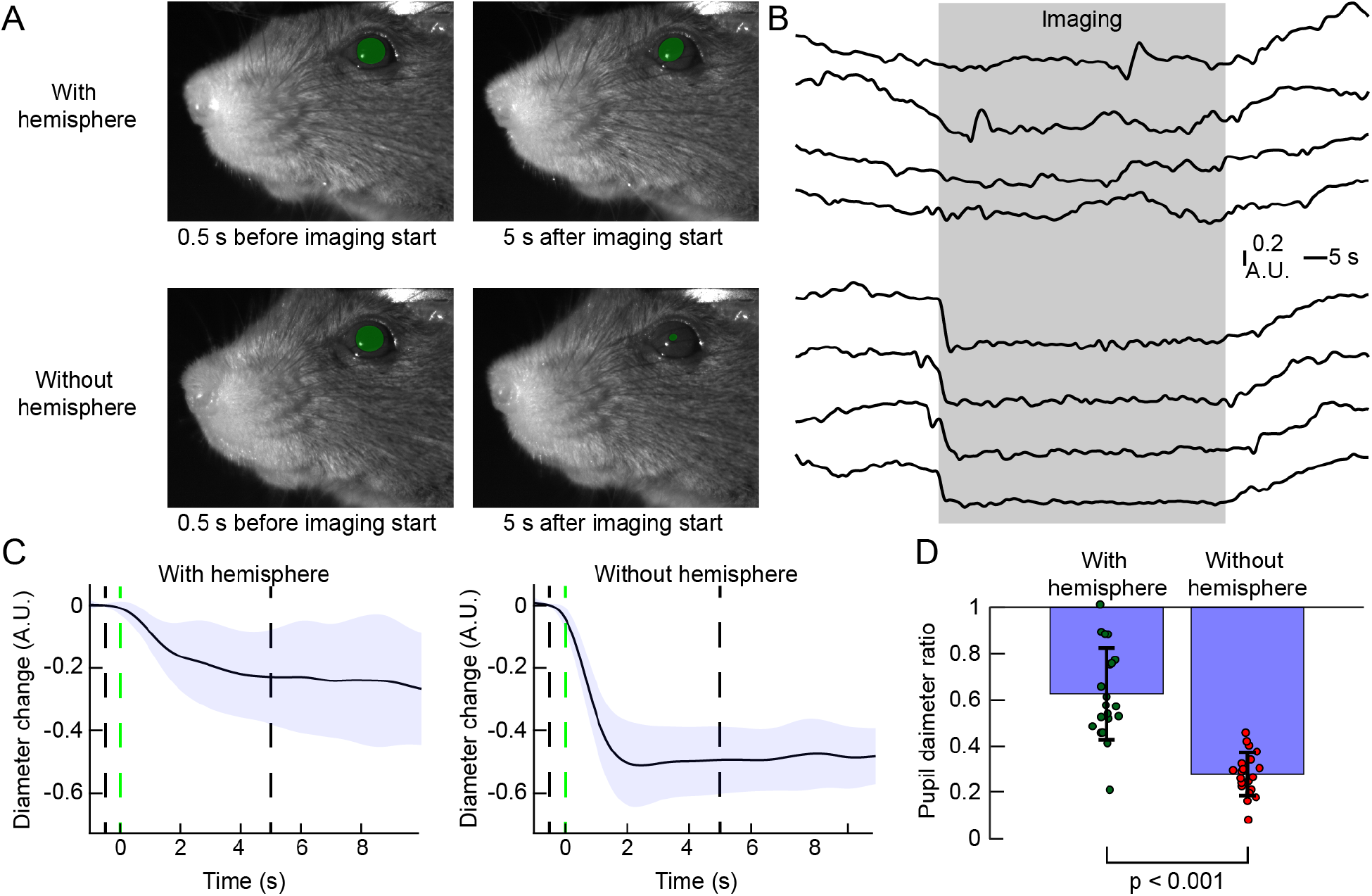
The aluminum hemisphere shields illumination and reduces pupil constriction during mesoscale imaging. **A**. Images of the mouse face 0.5 s before and 5 s after widefield imaging acquisition begins. Green circles highlight the pupils. **B**. Example two-minute pupil diameter time courses showing the off-to-on and on-to-off transition. The grey-shaded area indicates the period when widefield acquisition was performed **C**. Average pupil diameter changes upon beginning of widefield imaging (20 trials; traces show average ± standard deviation). Pupil diameter 1.5-0.5 s before imaging was defined as baseline for each trial. The green-dashed line indicates the start of widefield imaging, the black dashed lines indicate time points of the images shown in panel A. **D**. Comparison of the ratio of pupil size 2-3 s after start of imaging to pupil size 1.5-0.5 s before imaging. Significance was tested with a student’s t test.

## Discussion

Here, we describe a new widefield microscope design for mesoscopic imaging of dorsal mouse cortex. Several groups describe widefield calcium imaging with a single fluorescent indicator and concurrent hemodynamic imaging, while other groups describe two-color fluorescence widefield imaging without hemodynamic imaging, while we identified one report describing a system for combined two-color fluorescence and hemodynamic imaging using two cameras (Wang et al. 2022). Compared to previous publications, our system has several unique features:

First, the system enables quasi-simultaneous imaging of two spectrally distinct fluorophores (here: EGFP-based GRAB_ACh3.0_ and mApple-based jRGECO1a) together with quantitative estimates of Δ[HbO] and Δ[HbR] via two-color absorption imaging on a single detector. Changes in [HbO] and [HbR] are used to correct hemodynamic artifacts of the recorded fluorescence signals. Further, we can use them to study the relationship between cortical hemodynamics and different aspects of brain activity, such as neuronal firing and neuromodulation thereby gaining further insights into brain state-dependent regulation of cerebral blood flow and metabolism (Machler et al. 2021).

Second, we use a single detector. Using one camera drastically reduces the overall cost to build the system but requires sequential image acquisition for every wavelength (here: four wavelengths). We achieve sufficient SNR with negligible spectral crosstalk at a 10 Hz acquisition rate, which is within the range for widefield imaging studies in awake animals (Lake et al. 2020, Lohani et al. 2022).

Finally, the illumination light is contained within an aluminum hemisphere. Besides providing diffuse illumination for light from 525- and 625-nm LEDs to minimize reflections at the glass window, the hemisphere blocks most of the illumination light from the animal’s view. This prevents interference with behavioral measurements of pupil diameter as well as unwanted stimulation of the visual system. Pupil size measurements are often used as a non-invasive, indirect readout of alertness (Joshi et al. 2016, Reimer et al. 2016). We expect our shielded illumination design will enable seamless integration of mesoscale imaging with behavioral experiments based on visual stimulation (Vinck, Batista-Brito et al. 2015, Peters, Fabre et al. 2021) and virtual reality (Pinto, Rajan et al. 2019).

The overall cost to build our widefield imaging system with the components used is around $45,000, with the camera comprising the bulk of the cost (∼$25,000). While this is a lower price than for typical commercially available widefield imaging systems, modifications could further reduce the overall cost of the system. Depending on the brightness of the used fluorophores, their expression level, as well as experimental demands, sufficient imaging performance could be achieved with a less expensive camera (see **Supplementary Figure 6**). Furthermore, we used commercially available LED drivers as well as commercially available LEDs for fluorescence excitation. Using a custom design or already available open-source components with similar performance characteristics, especially regarding their response time to digital triggers, could further reduce the overall cost (see, e.g., (Harrison et al. 2009)).

Our imaging system can easily be modified or expanded to suit the requirements of different experimental studies; changing a LED and the corresponding optical filters will allow combining other fluorescent indicators if they are spectrally separated. This would, for example, allow integration of calcium indicators excited with near-infrared light (Qian et al. 2020). As the spectral diversity of genetically encoded molecular probes expands, widefield imaging with three spectrally separated fluorescent indicators might soon become a feasible approach. Furthermore, we and others combined mesoscale imaging with extracellular electrophysiology, providing simultaneous access to cortical network activity as well as single-or multi-unit activity in targeted cortical and subcortical areas (Allen et al. 2017, Thunemann et al. 2018, Lu et al. 2023).

Our current system design has the following limitations. First, despite aiming to minimize specular reflections and illumination inhomogeneity of 530- and 630-nm LEDs through diffuse illumination, we still observe inhomogeneous levels of absolute image brightness across the field of view. In practice, however, some variation of absolute brightness will not affect the readout of relative (i.e., intensity-normalized) changes in fluorescence or absorption, except for potentially reduced signal-to-noise levels in darker regions of the image. A possible way to further improve illumination homogeneity of both reflectance and fluorescence illumination is the incorporation of an integration device with uniformity correction, such as EUCLID (Celebi et al. 2023) into our imaging system. Further, uneven signal intensity in the medial-to-lateral direction is due to the curvature of the glass window covering the cranial exposure (see **Figure 2A**). Use of curved glass also causes different cortical depths to match the focal plane of the imaging system, which potentially introduces spatially heterogenous sampling bias. Here, use of adaptive lens optics, such as meta-lenses (Yang et al. 2023) could improve uniformity of illumination and could be used to match the focal plane across the field of view. Designing an imaging system with a single camera reduces overall system cost. However, the use of two cameras would allow for higher acquisition rates through spectral multiplexing. Further, the dual-band dichroic mirror and emission filter cut the available spectral bandwidth of the EGFP/GRAB_ACh3.0_ emission band (500-540 nm) and the mApple/jRGECO1a excitation band (550-570 nm). This potentially enables crosstalk due to off-peak excitation of the red fluorophore (see further discussion in **Appendix B**). However, with the fluorophores at the expression levels used, we achieve sufficient SNR with negligible spectral crosstalk at a 10-Hz acquisition rate, which is within the range of appropriate temporal resolution for widefield imaging studies in awake animals (Lake et al. 2020, Lohani et al. 2022). Furthermore, the extension of the imaging system by a second camera requires minimal modifications to the overall layout, allowing for higher acquisition speeds, if desired. Another consideration during the design of our system was the use of lasers instead of LEDs for excitation of the green and red fluorescent proteins. While less expensive, LEDs have a broad emission spectrum that requires clean-up through excitation filters. This limits the emitted light to the usable spectral range (see, e.g., **Figure 1E**, 565-nm LED) leading to an overall reduction in the available power compared to the nominal output (75 mW versus 8.6 W for the 565-nm LED), potentially requiring longer integration times to achieve sufficient SNRs. Here, use of a laser would enable excitation at higher power levels, if required. A drawback of using lasers instead of LEDs would be a substantial increase in the cost of the system.

In summary, given the versatility of our system and the comparatively low cost in contrast to commercial intravital imaging setups, we expect our widefield imaging system design to become a useful platform for neurovascular studies in the neuroscience and neurophotonics community.

## Methods

### Experimental animals

All procedures were conducted with approval from Boston University Institutional Animal Care and Use Committee and in accordance with the National Institutes of Health Guide for the Care and Use of Laboratory Animals. Standard rodent chow and water were provided ad libitum. Mice were housed in a 12-hour light cycle with lights turning on at 7:30 AM. We used three female mice of the Thy1-jRGECO1a GP8.20 (Dana et al. 2016, Dana et al. 2019) line; cortex-wide GRAB_ACh3.0_ expression was induced by injection of 1.5 µL AAV9-hSyn-GRAB_ACh3.0_ (WZ Biosciences, 3.06×10^13^ GC/mL) into each transverse sinus (3 µL per animal) of 1-day old neonates according to procedures described in Lohani et al. (2022).

Headpost implantation and craniotomy in adult (8-16 week old) animals follows procedures described in Kilic et al. (2020). Briefly, a custom-designed headpost machined from titanium was attached to the cranium and a modified crystal skull curved glass window was used to replace the dorsal cranium on both hemispheres (Kim et al. 2016). Prior to surgery, the original curved glass (12 mm width, labmaker.org) was cut in half to obtain separate glass pieces for each hemisphere. Following the implantation procedure, a silicone plug was placed on top of the glass window and a 3D-printed cap was fixed to the headpost to prevent heat loss. Mice were allowed to recover for at least one week following surgery before behavioral training commenced.

To adapt mice to be comfortable under head fixation, animals were handled until they remained calm in the gloved hand of the investigator. Mice were then head-fixed for increasing periods while they received a reward of sweetened condensed milk. Data acquisition was started when animals tolerated one hour of head fixation.

### Imaging system optical components

The imaging system is assembled from standard optomechanical components compatible with two-inch lens tubes. The imaging system is placed on an elevated platform (61×61 cm, Thorlabs, MB2424) on a vibration-isolating table.

A 470-nm LED (Thorlabs, SOLIS-470C) is used to excite GRAB_ACh3.0_ and a 565-nm LED (Thorlabs, SOLIS-565C) is used to excite jRGECO1a. A 466/40 nm excitation filter (Semrock, FF01-466/40, diameter 50 mm) is placed in front of the 470-nm LED and a 560/14 nm excitation filter (Semrock, FF01-560/14, diameter 50 mm) is placed in front of the 565-nm LED. A dichroic mirror combines excitation light from 565-nm and 470-nm LEDs (Semrock, FF520-Di02, 50 x 72 mm). The output of these LEDs is filtered through a 450-nm longpass filter (Chroma, AT450lp) to remove low wavelength light from the 565-nm LED that leaks through the primary excitation filter. A 45-degree mirror (Thorlabs, PF20-03-P01, diameter 50.8 mm) reflects excitation light down towards the mouse and reflects emitted light into the plane of the camera. A dual-band ‘main dichroic’ beam splitter (Semrock, Di03-R488/461, 50 x 72 mm) reflects excitation light towards the mirror and transmits emitted light towards the camera. A dual-band emission filter (Semrock, FF01-523/610, diameter 50 mm) is placed between a dichroic mirror and camera. A tube lens (Olympus, MVX-TLU-MVX10) projects the image onto the sCMOS camera (Andor, Zyla 4.2P-CL10 or Sona 4.2B-6).

An objective lens (Olympus, ZDM-1-MVX063) focuses the fluorescence excitation light onto the brain for epi-illumination and projects the emitted light towards the camera. The objective lens is surrounded by the reflective illumination ring (RIR) assembly, which contains six equally spaced 525-nm LEDs (CREE LED, XBDGRN-00-0000-000000C02) and six equally spaced 625-nm LEDs (CREE LED, XBDRED-00-0000-000000701) for reflectance imaging. Bandpass filters were cut to 9.4-mm diameter and placed in front of the reflectance LEDs. These filters are 532/10 nm bandpass filters (Edmund Optics, 65094) for the 525-nm LEDs and 632/10 nm bandpass filters (Edmund Optics, 65105) for the 625-nm LEDs. Six Peltier elements (Ferro Tec, 81044) controlled by a feedback-controlled temperature controller (Thorlabs, TED200C) actively cool the LEDs in the RIR to maintain the LEDs at room temperature during data acquisition. Temperature is measured on the LED ring with a 10-kΩ thermistor (TDK, NTCG163JF103FT1). Heat from the Peltier elements is transferred to a custom heat sink made of aluminum. The RIR allows fixation of custom hemisphere made of aluminum (bead-blasted) with a 15-mm opening located directly above the cranial window.

To perform imaging tests with a different camera model (ac1920-150uc, Basler with Python 2000 CMOS chip from ON semiconductor in global shutter mode), a 45-degree mirror (Edmund Optics, 83-538) behind the detection filter was used to divert the light path to a second camera arm with alternative tube lens (ThorLabs, TTL180-A). The camera was controlled through the manufacturer’s software (Pylon Viewer); 4x4 binning (sum) with analog gain disabled were used with exposure times set to 6 ms for all frames.

### Control of imaging acquisition

We use a computer-controlled DAQ device (National Instruments, PCIe-6363) to operate the imaging system. The DAQ device is connected to two I/O connector boards (National Instruments, CB-68LPR) inside a control interface box. The control interface relays a DAQ-generated exposure trigger to the camera. An integrated circuit with an OR gate (Texas Instruments, SN74AHC32N) within the control interface receives the “Fire ALL” trigger from the camera as an input. The “Fire ALL” trigger is positive when the exposure state overlaps for all rows in the rolling shutter mode. The output of the OR gate is the common input to an integrated circuit with four AND gates (Texas Instruments, SN74AC08N). The second – wavelength-specific – input to each individual AND gate is the LED selection trigger generated by the DAQ system. Each output of the AND gate connects to one LED driver (Thorlabs, DC2200) controlling one type of LED. During normal operation of the imaging system, the “Fire ALL” trigger controls when the LED turns on. The other input to the OR gate is a camera bypass trigger that allows the LEDs to be triggered while the camera is off. We use a custom MATLAB script to control the DAQ system. We used image capture software (Andor, SOLIS) to control camera and image acquisition.

The animal was positioned under 525-nm illumination and the platform carrying the animal was adjusted to bring the brain surface into focus. Then, the aluminum dome is lowered to minimize the distance between its opening and the brain surface. Illumination intensities (i.e., LED currents) were adjusted to bring image intensities into the dynamic range of the camera at 16-bit image depth. The acquisition window was cropped to the size of the field of view; image height (number of rows) was a variable entered into the MATLAB script controlling image acquisition: the length of the period when all rows are exposed (“FIRE all”) depends on image height and requires adjustment of the camera exposure trigger generated by the DAQ system. Typical image sizes were 460×360 pixels with a 9.2×7.2 mm field of view and 20-μm resolution at 4×4 binning. Based on selected exposure times and frame rate, the MATLAB script calculates the total number of frames per acquisition run; this number is entered into the image capture software. The camera is set to “External Exposure” mode and image acquisition is started. Then, execution of the MATLAB script is started causing the DAQ interface to send LED selection and camera triggers to the control interface.

### Sensory stimulation and behavioral recordings

The MATLAB acquisition script generates additional triggers for synchronization with behavioral apparatuses, such as to deliver stimuli by puffing compressed air to the animals’ whiskers. Stimuli are defined by duration, frequency, repetitions, interstimulus interval for individual stimuli and sequences of pre-stimulus baseline, stimulus, and post-stimulus observation period. Based on these parameters, the MATLAB script calculates the required imaging time and number of images. For air puff stimuli, the 5-V trigger is sent to a picopump (WPI, PV830) connected to a 2-mm diameter plastic tube that terminates in a glass capillary to deliver pressurized air to the mouse whiskers.

The animal’s face and whiskers are recorded during imaging with a CMOS camera (Basler, acA1920-150uc) through a variable zoom lens (Edmund Optics, 67715). A 940-nm LED (Thorlabs, M940L3) continuously illuminates the animal, and a 920-nm long pass filter prevents light from the microscope from contaminating the camera recording. The mesoscope control system triggers the CCD camera to acquire a single frame during each acquisition cycle.

An accelerometer (Analog Devices, ADXL335) is placed underneath the mouse bed to record animal’s motion. The accelerometer generates analog signals for motion in X, Y and Z direction which are recorded by a separate DAQ system (NI, USB-6363) receiving a synchronization signal from the mesoscope control system.

### Image processing

The camera control software (Solis, Andor) stores the acquired images as raw data (‘.dat’) together with metadata files (‘.ini’ and ‘.sifx’). For all subsequent import and processing steps, we use MATLAB. Raw image data is loaded into MATLAB, bypassing conversion into image (.tif) files by the camera software and enabling multi-stream data import from high-performance solid-state drives with MATLAB’s multi-processor environment. For all channels, a baseline image representing the temporal average of all images for that channel is generated. For fluorescence channels, ΔF/F images are computed on a pixel-by-pixel basis by dividing each image by the baseline image. After calculating ΔF/F, linear trends are removed from the time course of each pixel using the MATLAB function “detrend”. The modified Beer-Lambert law is used to calculate changes in HbO and HbR concentrations from 525 nm and 625 nm reflectance (see implementation for MATLAB’s multi-processor environment in **Appendix A**). Estimated Δ[HbO] and Δ [HbR] are used to correct green fluorescence measurements for artificial intensity changes caused by dynamic changes in excitation and emission light absorption by hemoglobin. The correction method has been described in Ma et al. (2016a) and its implementation is further detailed in **Appendix A**. In this correction approach, the measured fluorescence is equal to the artifact-free change in fluorescence multiplied by a correction factor containing the pathlength in tissue and the tissue absorption coefficient. Pathlengths have been previously estimated based on Monte-Carlo simulations (Ma et al 2016). Since hemoglobin accounts for most of the light absorption in brain tissue, light absorption depends largely on Δ[HbO] and Δ[HbR] and the known HbO and HbR extinction coefficients at the respective wavelengths of fluorescence excitation and emission. We calculate the light absorption coefficient for every pixel at every time point based on our estimations of Δ[HbO] and Δ[HbR] and apply it to correct for hemodynamic darkening of ΔF/F in the green (EGFP/GRAB_ACh3.0_) channel.

Measurements with several consecutive trials are interpolated at 100-ms intervals using a time vector relative to the stimulus onset. Trial-averaged ratio maps are calculated by averaging individual time courses across trials.

Behavioral video recordings are stored as image sequences and imported into MATLAB. To calculate pupil size, an elliptical region is drawn defining the eye and an intensity threshold is manually defined. The pupil size is calculated as the number of pixels below the threshold. The resulting time series is normalized to its maximal and minimal pupil size within the recording.

#### *In vitro* crosstalk calibration

Solutions of purified 0.03 mg/mL EGFP (Abcam, ab84191) and 0.4 mg/mL mApple (kindly provided by Ahmed Abdelfattah) in phosphate-buffered saline (PBS) were prepared prior to imaging. 30-μL samples of EGFP, mApple, or PBS-only control solution were each injected into one channel of separate 6-channel slides (ibidi, µ-Slide VI – Flat). The center of the fluorophore-filled channel was focused under the widefield imaging system before starting image acquisition for each slide. Illumination during an imaging sequence alternated between the 470-nm and 565-nm LEDs while stepping through a series of camera exposure times and increasing LED currents. Three trials were conducted for each sample, with the position of the slide under the objective being shifted between trials to account for intensity variations. To calculate the fluorescence intensity, pixel values were averaged over a region of interest covering the center of the fluorophore-containing channel. Then, we took the mean of the three trials per condition and subtracted the fluorescence intensity of the PBS control. Further analysis of crosstalk is detailed in **Appendix B**.

### Photo-switching of jRGECO1a

A variant of the MATLAB acquisition script was used to create two types of acquisition cycles: One cycle includes 470-nm illumination and disables image acquisition (no camera exposure trigger and bypassing the ‘FIRE ALL’ trigger), while the other cycle does not include 470-nm illumination but normal image acquisition. The combination of both cycles enables intermittent 470-nm illumination cycles at variable frequencies. Image analysis is performed on baseline-normalized time courses averaged from all pixels across the brain surface.

## Acknowledgements

We thank Ahmed Abdelfattah for generously providing purified mApple fluorescent protein for calibration of our imaging system. We gratefully acknowledge support from the NIH (BRAIN Initiative R01MH111359, BRAIN Initiative U19NS123717, R01DA050159, R01NS108472). Patrick Doran was supported by the Ruth L. Kirschstein Predoctoral Fellowship F31NS118949. Rockwell P. Tang was supported by NIGMS T32GM145455. The authors acknowledge that data analysis was performed on the Shared Computing Cluster which is administered by Boston University’s Research Computing Services. We thank the members of the Neurovascular Imaging Laboratory for helpful discussions.

## Data availability

Design blueprints, parts inventory, data acquisition and analysis software are available on github (https://github.com/NIL-NeuroScience/WidefieldImagingAnalysis); raw and processed data are available on Zenodo (for Figure 2, resting state: https://doi.org/10.5281/zenodo.10798934; for Figure 2 stimulus: https://doi.org/10.5281/zenodo.10798996; for Figure 3: https://doi.org/10.5281/zenodo.10798658; for Supplementary Figure 3: https://doi.org/10.5281/zenodo.10798091; for Supplementary Figure 6 resting state: https://doi.org/10.5281/zenodo.10805179; for Supplementary Figure 6 stimulus: https://doi.org/10.5281/zenodo.10805381).

## Author contributions

Patrick R. Doran: Investigation, Formal Analysis, Software, Visualization, Writing – original draft, Writing – review & editing; Natalie Fomin-Thunemann: Investigation, Writing – review & editing; Rockwell P. Tang: Investigation, Visualization, Writing – original draft, Writing – review & editing; Dora Balog: Software, Writing – review & editing; Bernhard Zimmermann, Kıvılcım Kılıç, Joel D. Herbert, Grace Chabbott, Bradley C. Rauscher, John X. Jiang: Resources, Writing – review & editing; Sreekanth Kura, Harrison P. Fisher: Software, Writing – review & editing; Sava Sakadzic, David A. Boas: Conceptualization, Writing – review & editing; Anna Devor: Conceptualization, Funding acquisition, Supervision, Writing – original draft, Writing – review & editing; Ichun Anderson Chen: Conceptualization, Methodology, Supervision, Writing – review & editing; Martin Thunemann: Conceptualization, Software, Supervision, Visualization, Writing – original draft, Writing – review & editing.

## Appendix

## Appendix A: Optimization of Δ[HbO] and Δ[HbR] estimation process for MATLAB using an explicit solution of HbR and HbO with measured reflectance at 525 nm and 625 nm

Concentration changes of deoxy- and oxyhemoglobin ([*HbR*] and [*Hbo*]) can be estimated using a system of linear equations (here, we use intensity measurements at two wavelengths (525 nm and 625 nm) and two unknowns ([*HbR*] and [*Hbo*]). For further discussion, see Ma et al. (2016).

The molar extinction coefficients *ε* of Hb and HbO at 525 nm and 625 nm are known variables.

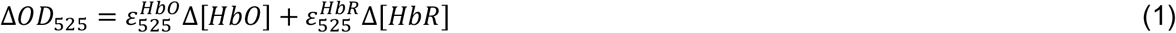

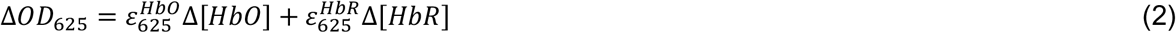

We estimate the optical density in every pixel for every time point t from the measured intensities 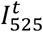 and 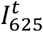 as follows:

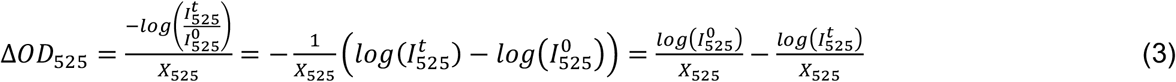

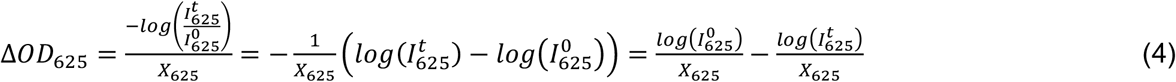

The pathlengths *X*_525_ and *X*_625_ are known variables; the “baseline” intensities 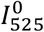 and 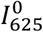 are estimated using the temporal average of the intensities at the given pixel.

When we solve the equations (1) and (2) explicitly, combine them with equations (3) and (4), and combine all variables that are constant over space and/or time, we enable efficient (i.e., fast) approach for estimating [*Hb*] and [*Hbo*].

To estimate Δ[*Hbo*], we first solve equations 1 and 2 for Δ[*Hb*]:

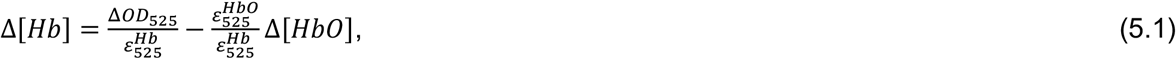

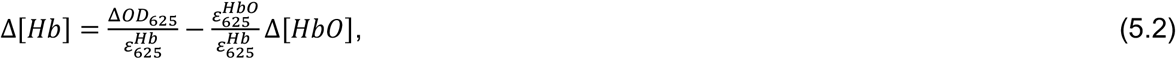

and combine equations 5.1 and 5.2:

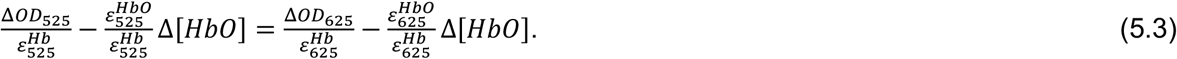

Then, we solve equation 5.3 for Δ[*Hbo*]:

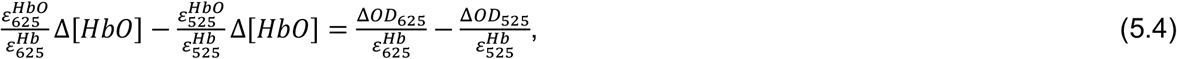

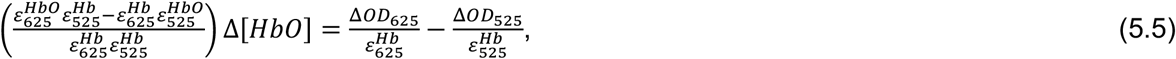

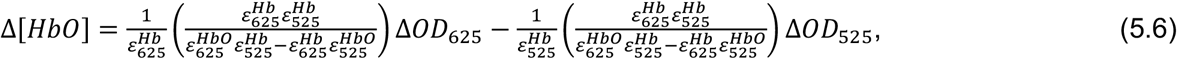

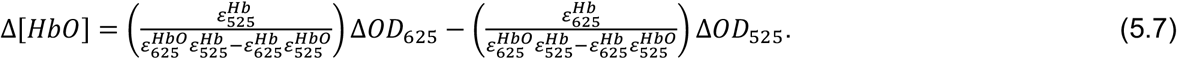

Then, we combine all constant variables in equation 5.7 to define 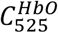 and 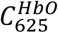:

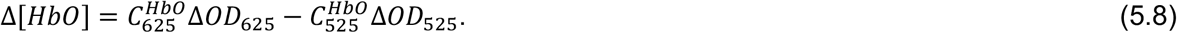

Now, we insert the definition of Δ*OD*525 and Δ*OD*625 from equations 3 and 4:

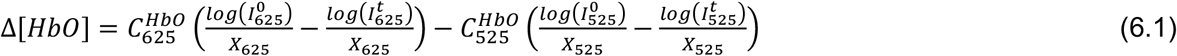

and combine again all variables that are constant over time:

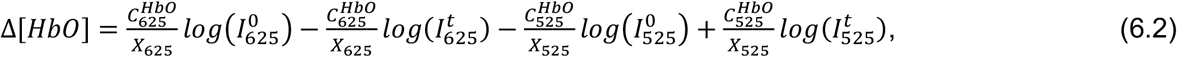

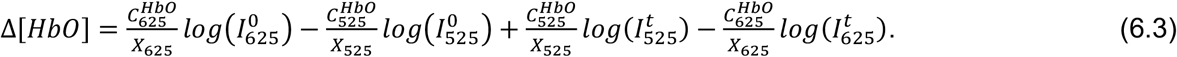

Finally, we combine all constant variables into three parameters 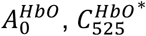, and 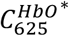:

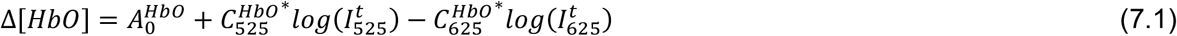

With

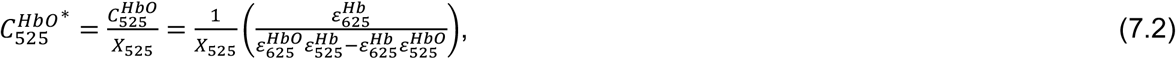

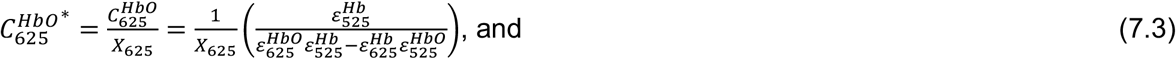

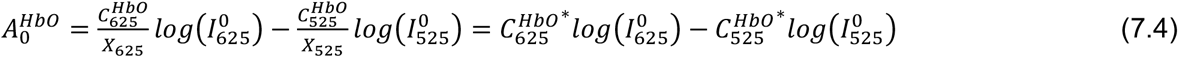

*Note that* 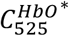*and* 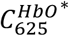 *are invariant in time and space, while* 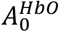 *is only time-invariant – its value changes for every pixel*.

In the same way, we can estimate [*Hb*]. First, we solve equations 1 and 2 for [*Hbo*]:

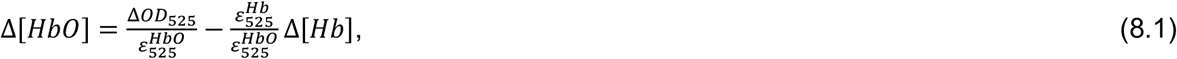

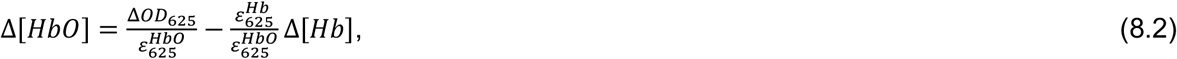

and combine equations 8.1 and 8.2:

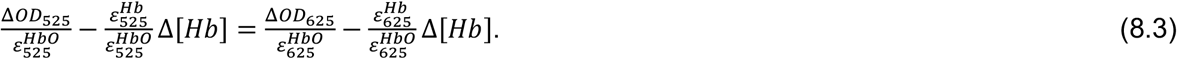

We solve equation 8.3 for Δ[Hb]:

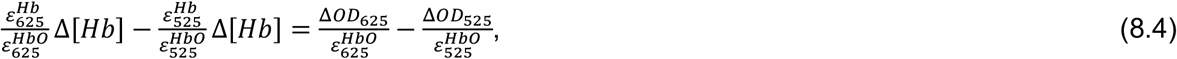

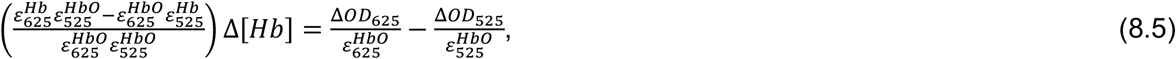

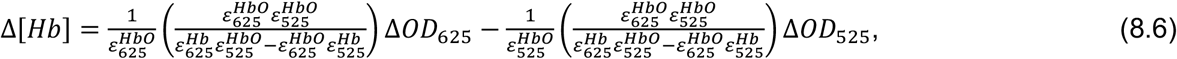

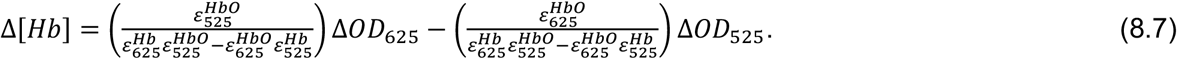

We combine all constant variables in equation 8.7 to define 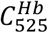 and 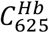

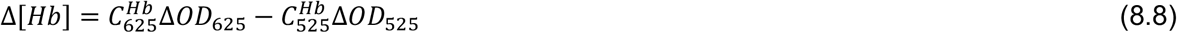

and insert the definition of Δ*OD*525 and Δ*OD*625 from equations 3 and 4:

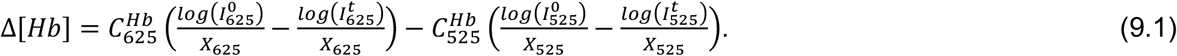

Then, we combine again all variables that are constant over time:

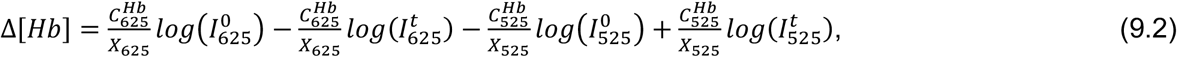

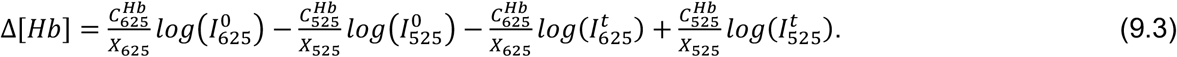

Finally, we combine all constant variables into three parameters 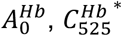, and 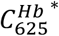:

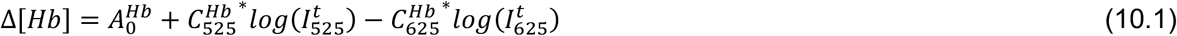

With

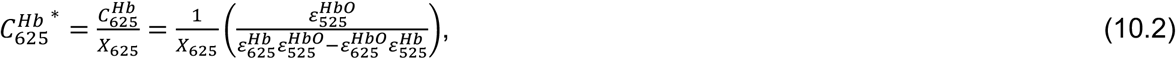

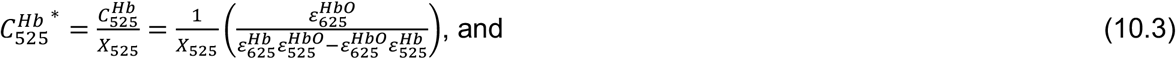

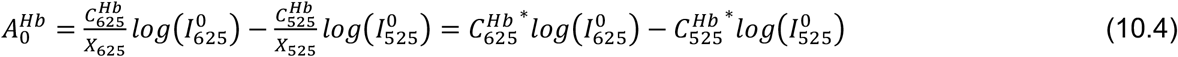

*Note that* 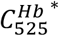 *and* 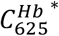 *are invariant in time and space, while* 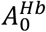 *is only time-invariant – its value changes for every pixel*.

*MATLAB implementation:*

(1) Calculate 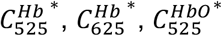, and 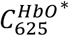 [dim: 1]

(2) Calculate 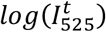 and 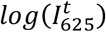 [dim: WxHxT]

(3) Calculate 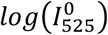 and 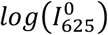 [dim: WxH]

(4) Calculate 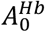 and 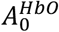 [dim: WxH]

(5) Estimate 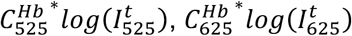 [dim: WxHxT]

(6) Combine 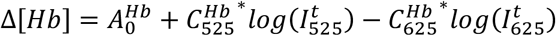 [dim: WxHxT]

(7) Estimate 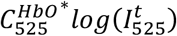, and 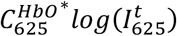 [dim: WxHxT]

(8) Combine 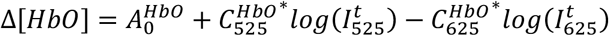 [dim: WxHxT]

## Appendix B: Evaluation of potential crosstalk between EGFP- and mApple-derived fluorophores

### Part I: in vitro calibration with purified EGFP and mApple fluorescent proteins

We performed fluorescence measurements of purified EGFP and mApple that were excited at 475 nm and 565 nm using series of increasing exposure times and optical powers. For further evaluation, we calculated the radiant energy *E* for each measurement as the product of illumination power, estimated from the LED current using a separately measured power calibration curve, and exposure time.

Then, we defined the illumination ratio *R* for EGFP and mApple as radiant energy at off-peak illumination divided by radiant energy at on-peak illumination combining different illumination conditions for on- and off-peak excitation into a single variable:

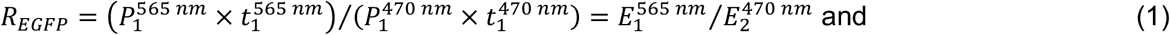

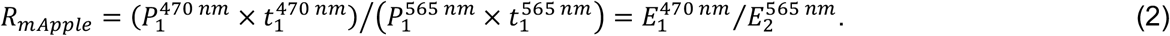

Note that every illumination ratio can be achieved by different combinations of optical powers and exposure times. This means that pairs of fluorescence measurements are performed with different combinations of optical power and exposure time for off- and on-peak excitation but match the chosen illumination ratio. The corresponding individual on- and off-peak fluorescence intensities have different values. We define their ratio, measured at a certain LED power *P*1 and exposure time *t*1 at off-peak excitation relative to the fluorescence intensity at a certain LED power *P*2 and exposure time *t*2 at on-peak excitation for each fluorophore as crosstalk ratio C for EGFP and mApple as

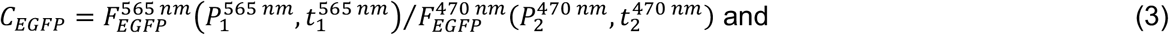

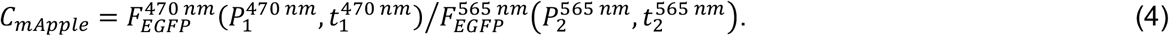

We again substitute LED power and exposure time by radiant energy *E* :

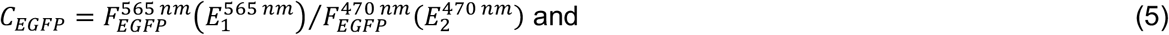

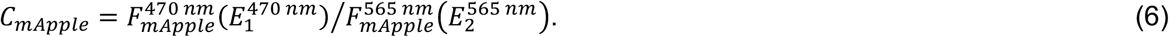

To estimate the crosstalk ratio C, we chose 20 pairs of measurements matching a given illumination ratio *R*. The pairs of fluorescence intensities from off-peak excitation versus on-peak excitation, when plotted against each other, follow a linear relationship (**Supplementary Figure 2C, D**), which is expected given the underlying linear relationship between fluorescence intensity and radiant energy (shown in **Supplementary Figure 2A, B**). Accordingly, we can estimate the crosstalk ratio C from the slope of the linear fit between fluorescence from off- and on-peak excitation for the 20 pairs of values for each given illumination ratio (**Supplementary Figure 2C, D**). Finally, for each fluorophore we plot the crosstalk ratio C versus illumination ratio *R* for EGFPand mApple (i.e., *CEGFP* versus *E*565 *nm*/*E*470 *nm* and *CmApple* versus *E*470 *nm*/*E*565 *nm* ; **Supplementary Figure 2E**). In the resulting graph, a small value of *R* indicates weak off-peak excitation compared to on-peak excitation, and the resulting crosstalk ratio C is small. For increasing values of *R*, off-peak excitation becomes increasingly stronger compared to on-peak excitation leading to a higher crosstalk ratio C. Given that EGFP shows only very weak off-peak excitation at 565 nm, its crosstalk ratio C increases only modestly with R. In contrast, as mApple is more efficiently excited off-peak at 470 nm, the crosstalk ratio C rises faster with increasing *R*.

#### Part II: in vivo correction of jRGECO1a crosstalk upon 470-nm excitation in GRAB_ACh3.0_ and jRGECO1a-expressing animals

In experiments with jRGECO1a- and GRAB_ACh3.0_-expressing mice (**Figure 2**), the used LED powers and exposure times for fluorescence excitation at 470 and 565 nm define a range of illumination ratios. Based on the *in*-*vitro* calibration with EGFP and mApple, we estimated that for EGFP-derived GRAB_ACh3.0_, <1% of its fluorescence present at 470-nm excitation contributes to fluorescence measured at 565-nm excitation, while for mApple-derived jRGECO1a, 3-9% of its fluorescence present at 565-nm excitation contributes to fluorescence measured at 470-nm excitation (highlighted areas in **Supplementary Figure 2E)**.

Calibration of the crosstalk ratio C enables us to estimate the level of fluorescence present at 470-nm excitation due to off-peak excitation of jRGECO1a for a given combination of illumination power and exposure times, i.e., radiant energies *E*1 and *E*2 :

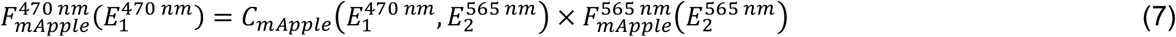

We assume here that fluorescence measured at 565 nm is solely due to jRGECO1a fluorescence with negligible contribution from GRAB_ACh3.0_, given the minor (<1%) contribution of EGFP fluorescence at 565 nm excitation. This enables us to define the following correction term to remove jRGECO1a crosstalk from fluorescence measured at 470-nm excitation:

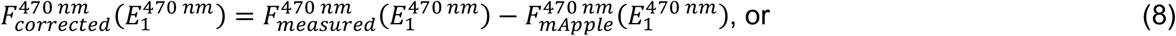

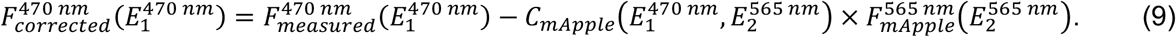

To apply this correction approach based on *in-vitro* determined crosstalk ratios *C* to *in-vivo* data, we make the following assumptions: (a) fluorescence acquired at 565 nm is solely caused by jRGECO1a and (b) dynamic changes in jRGECO1a fluorescence between consecutive acquisition of images at 470-nm and 565-nm excitation are negligible. Further, we do not consider wavelength-dependent absorption of light for fluorescence excitation and emission through hemoglobin. Incorporating [HbO] and [HbR] estimates into the correction would result in combining measurements from all four wavelengths to correct for crosstalk in fluorescence measurements at 470-nm excitation, potentially confounding this measurement, i.e., GRAB_ACh3.0_ fluorescence, by the sum of measurement noise from all measured channels.

To verify that the degree of in-vivo crosstalk matches our in-vitro estimation, following equation (9), we assume that we can estimate the “true” GRAB_ACh3.0_ signal by subtracting a proportion C of the 565-nm fluorescence image intensity from the 470-nm fluorescence image on a pixel-by-pixel and frame-by-frame basis. Here, we subtracted on a frame-by-frame three different crosstalk ratios (we tested C=0, 3%, 5%, 10%) and estimated the cross-correlation of the resulting, corrected F_470_ timeseries with the F_565_ timeseries (**Supplementary Figure 3**). For C of 5% and 10%, we observe clear, negative zero-lag cross-correlation indicative for overcorrection. This result suggests that the degree of crosstalk established under in vitro conditions is a reasonable estimate of the crosstalk we observe in vivo. Based on this evaluation, we suggest that under the illumination conditions for in vivo imaging used here, the benefits of spectral crosstalk correction are minimal. Therefore, we did not perform crosstalk correction for the data shown in **Figure 2**.

However, it is possible that levels of crosstalk increase, particularly when fluorescence from on-peak excitation of one fluorophore approaches levels of the other fluorophore due to off-peak excitation at the same wavelength. First, this could happen due to low expression of one fluorescent indicator versus the other. Here, using AAV delivery into newborn pups, we achieved relatively high levels of GRAB_ACh3.0_, diminishing the potential effect of jRGECO1a crosstalk on GRAB_ACh3.0_ measurements. Second, this situation could also arise when one fluorescent indicator has low levels of basal fluorescence in virtual absence of its analyte, i.e., acetylcholine for GRAB_ACh3.0_ in this study. When ACh levels are low (low GRAB_ACh3.0_ fluorescence) and neuronal calcium levels are high (high jRGECO1a fluorescence), the measurement at 470-nm excitation could potentially show higher contribution from jRGECO1a fluorescence compared to periods where ACh levels are high. To further address and quantify this effect, however, is very difficult with the used animals under the given imaging conditions as jRGECO1a and GRAB_ACh3.0_ signals show high correlation due to their underlying physiological covariance.

