## Supplementary Material for "Widefield *in vivo* imaging system with two fluorescence and two reflectance channels, a single sCMOS detector, and shielded illumination"

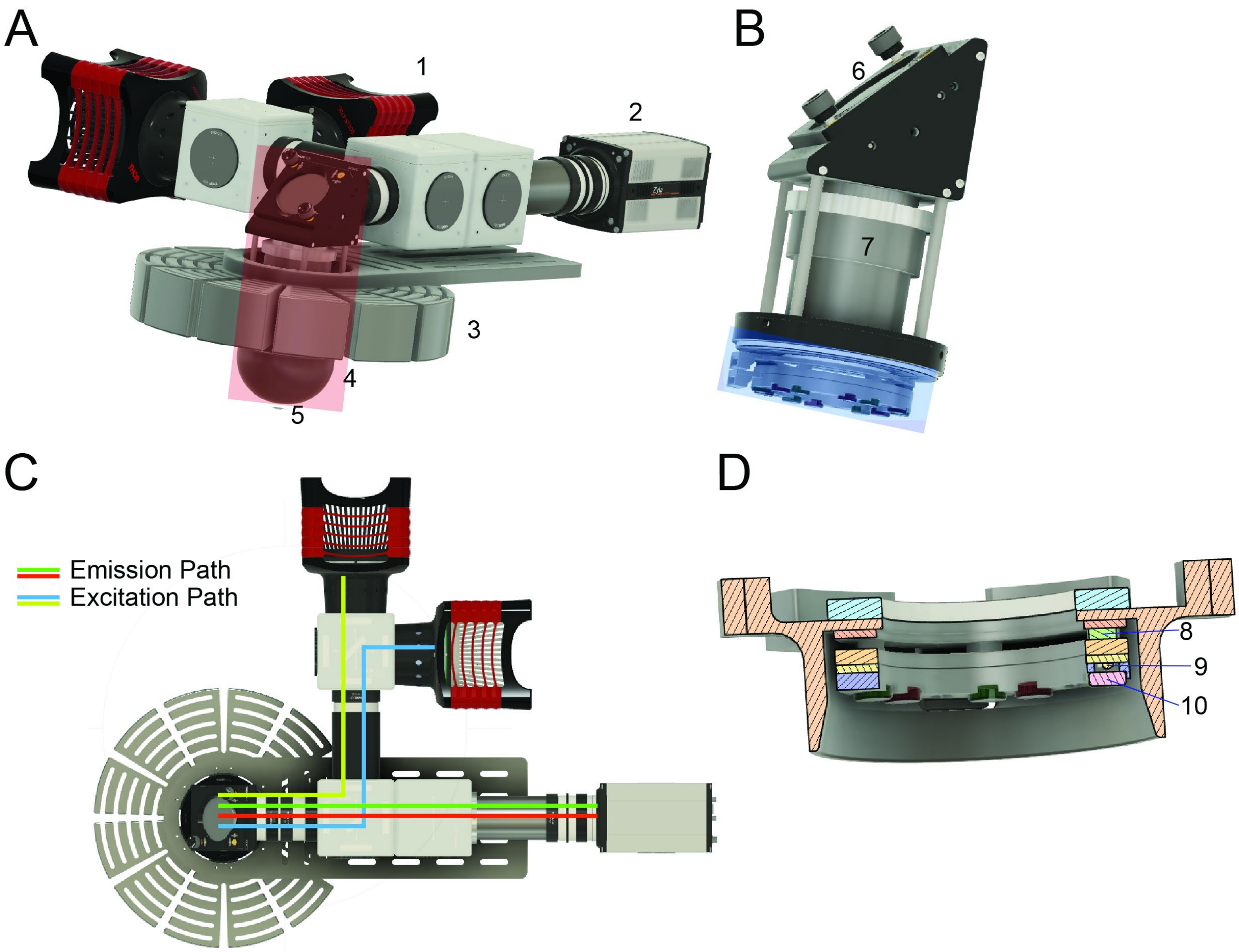


**Supplementary Figure 1: 3D model of the widefield imaging system. A.** Overview of the full imaging system showing (1) 470-nm LED, (2) sCMOS camera, (3) aluminum heat sink, (4) aluminum hemisphere, and (5) cranial window. The red rectangle indicates the area detailed in panel B. Designs of parts manufactured by Thorlabs were obtained from thorlabs.com and included here with permission of Thorlabs Inc. The design of the Andor Zyla camera was obtained from grabcad.com. **B.** This part of the system contains (6) 45° mirror and (7) objective lens. The blue rectangle highlights the reflectance illumination ring (RIR) detailed in panel D. **C**. Top view of the imaging system indicating the light paths of fluorescence excitation and emission. The light is reflected down from the plane shown in this panel into the objective by a 45° mirror (number 6 in panel B). **D**. Cross section of the RIR showing the arrangement of rings containing (8) Peltier elements, (9) 525- and 625 nm LEDs, and (10) excitation filters.

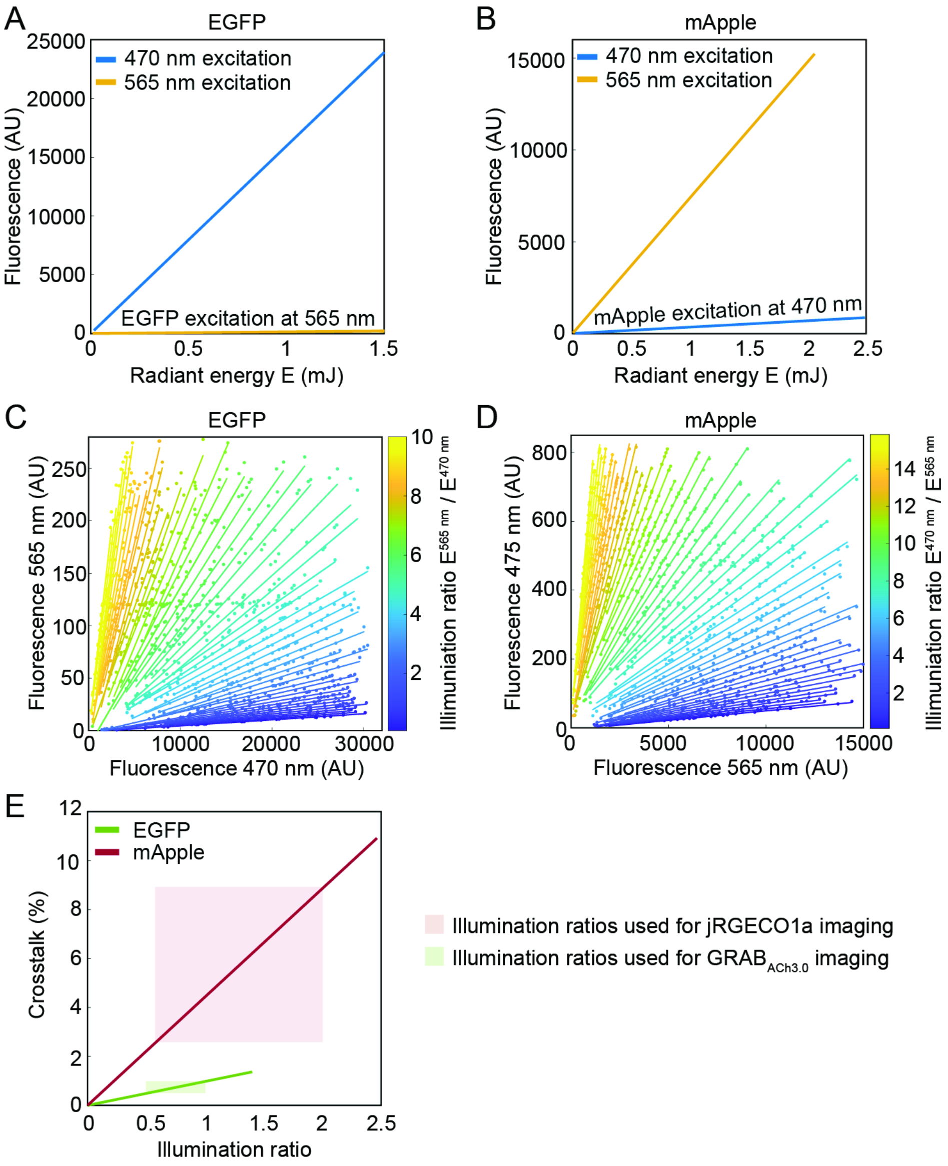


**Supplementary Figure 2: Estimation of fluorescence crosstalk using purified EGFP and mApple proteins.** See **Appendix B** for further details and discussion. **A, B.** EGFP (A) and mApple (B) fluorescence was measured using the widefield imaging system for various combinations of illumination power and exposure time for the 470-nm and 565-nm LEDs. Illumination power and exposure time were multiplied to estimate radiant energy. **C, D.** Relative fluorescence of EGFP (C) and mApple (D) at different illumination conditions for on- and off-peak excitation. Lines connect pairs of fluorescence measurements where on- and off-peak excitation have the same illumination ratio, i.e., where the ratio of off- versus on-peak radiant energy is the same. For each illumination ratio, twenty pairs of measurements with that ratio were chosen and the linear fit between on- and off-peak fluorescence defines the relative crosstalk for the given illumination ratio. **E.** Relative crosstalk due to off-peak excitation at different illumination ratios for EGFP and mApple. The transparent rectangles cover the range of illumination ratios used during in-vivo imaging with EGFP-derived $GRAB_{ACh3.0}$ and mApple-derived jRGECO1a.

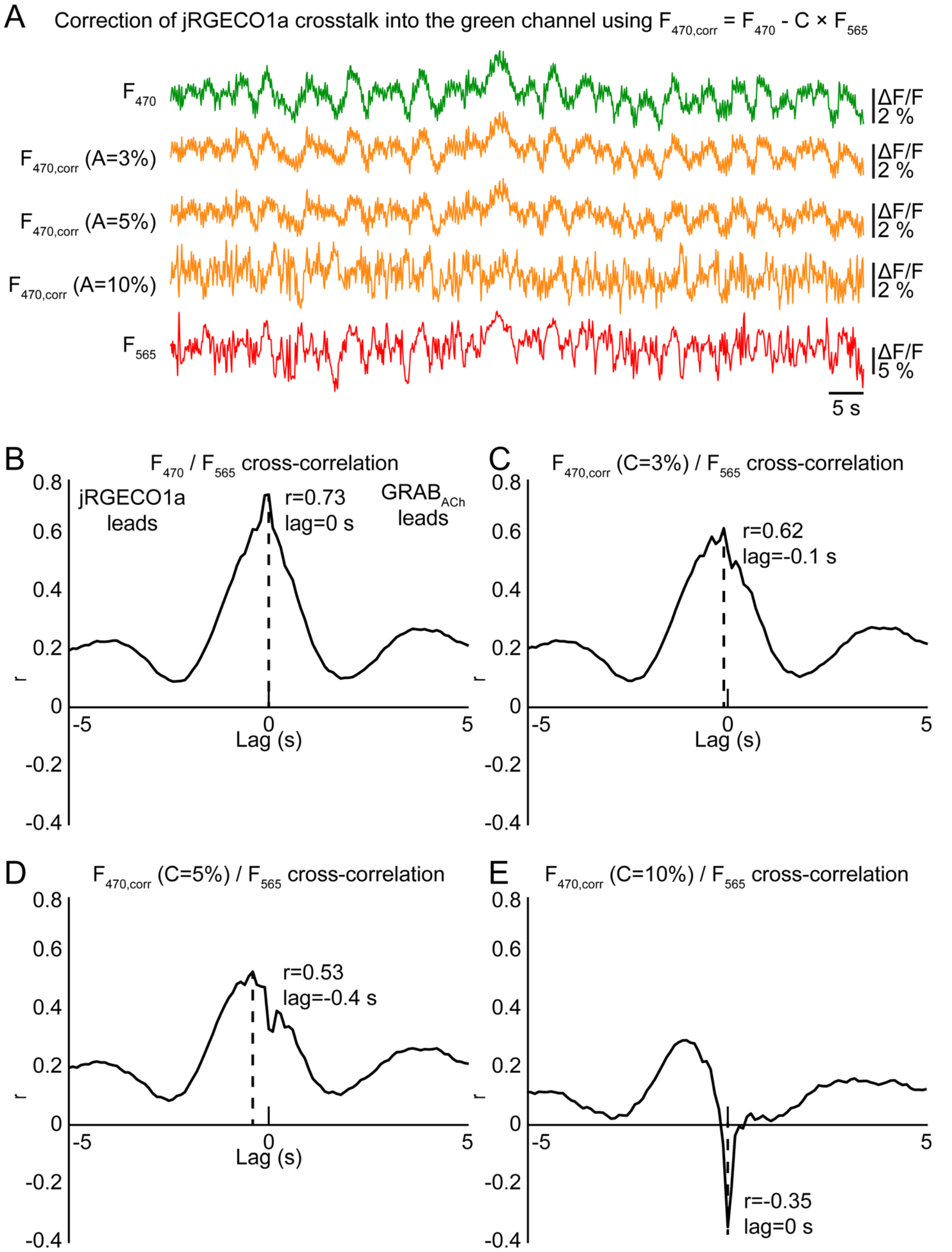


**Supplementary Figure 3. *In-vivo* characterization of jRGECO1a fluorescence crosstalk. A.** Traces show spontaneous activity recorded in the left barrel cortex of an awake, jRGECO1a- and $GRAB_{ACh3.0}$-expressing mouse with an illumination ratio of 0.68 (see Supplementary Figure 2). This illumination ratio is estimated to cause a contribution of C=3% from jRGECO1a fluorescence at 565-nm excitation to the signal obtained at 470-nm excitation. The contamination of $GRAB_{ACh3.0}$ fluorescence with jRGECO1a fluorescence upon 470-nm excitation was therefore removed on a pixel-by-pixel basis by subtracting the crosstalk fraction C times the fluorescence intensity at 565-nm excitation ($F_{565}$) from the fluorescence intensity at 470-nm excitation ($F_{470}$). The $F_{565}$ time course (considered solely representing jRGECO1a fluorescence) is shown in red, and the uncorrected $F_{470}$ time course is shown in green. $F_{470,corr}$ time courses with different degrees of correction (C=3%, 5%, or 10%) are shown in orange. **B.-E.** Cross-correlation between the $F_{565}$ (i.e., jRGECO1a) time course and (B), the uncorrected ($F_{470}$) time course; (C), the $F_{470,corr}$ time course with C=3%; (D), the $F_{470,corr}$ time course with C=5%; or (E), the corrected $F_{470,corr}$ time course with C=10%. Cross-correlation analysis is based on a 10-minute recording of spontaneous activity in the left barrel cortex. Note the occurrence of negative correlation at zero lag in panel D and E. For further discussion, see **Appendix B** in this Supplement.

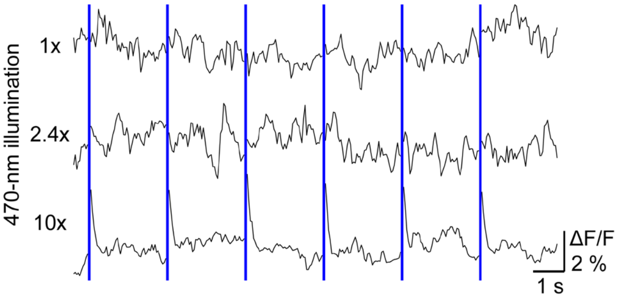


**Supplementary Figure 4. Photo-switching effect of on jRGECO1a upon 470-nm illumination.** Recording of jRGECO1a fluorescence in a Thy1-jRGECO1a transgenic mouse; traces represent intensity averaged over the cortical surface. Pulses of 470-nm light were delivered for 30 s at a rate of 0.2 Hz (blue bars); trial average of 10-20 trains of light pulses are shown. From top to bottom, traces show runs with increasing intensities of 470-nm illumination compared to typical intensities used during imaging in $GRAB_{ACh3.0}$- and jRGECO1a-expressing animals (1x, 79 mW with illumination time of 3 ms), 2.4-times the intensity (194 mW with illumination time of 3 ms), and ten times the intensity (304 mW with illumination time of 8 ms) used during imaging experiments. Frames acquired during 470-nm illumination showed artificial reduction in signal intensity and were omitted from the plots.

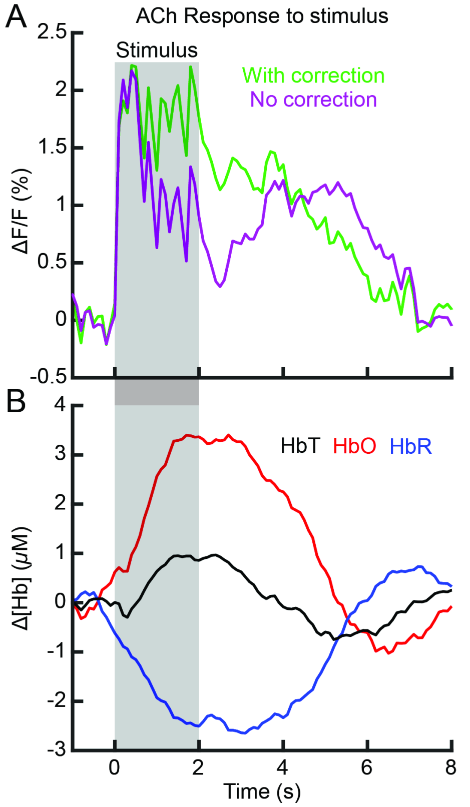


**Supplementary Figure 5: Correction of the hemodynamic artifact in GRAB$_{ACh3.0}$ fluorescence. A.** Trial average of stimulus-evoked ACh release in the contralateral barrel cortex in response to a 2-s train of air puffs at 3 Hz (20 trials). Time courses of GRAB$_{ACh3.0}$ fluorescence before and after hemodynamic correction are shown as purple and green lines, respectively. The data is the same as in Figure 2D. The grey shaded area indicates when the stimulus occurs. **B.** Time courses of oxy- and deoxyhemoglobin, and total hemoglobin (HbO, HbR, HbT; average of 20 trials) corresponding to the time course in panel A. The darkening of the uncorrected GRAB$_{ACh3.0}$ fluorescence time course (magenta line in panel A) is largest during the peak of the hemodynamic response around 1.5 - 3.5 s after stimulus onset.

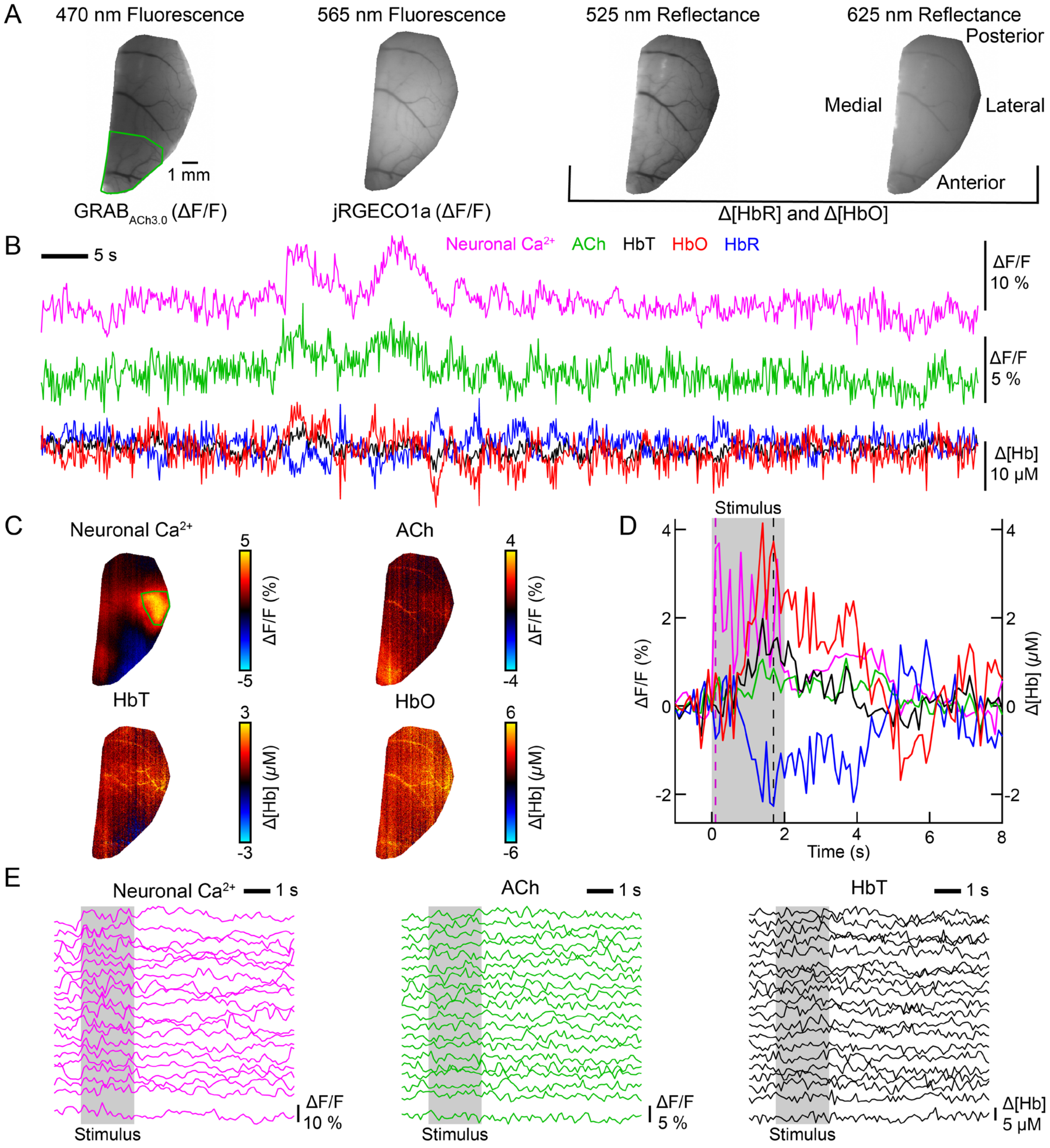


**Supplementary Figure 6**: Measurement of spontaneous and stimulus-evoked activity with an alternative (less expensive) camera at 4x4 binning and otherwise similar acquisition parameters as shown in Figure 3. **A**. Average of the first ten images collected during a ten-minute acquisition for each illumination wavelength. A mask was manually drawn around the cranial window. A green polygon was drawn around the secondary motor cortex. **B**. Spontaneous activity in the secondary motor cortex (green polygon in panel A) for calcium, acetylcholine (ACh), and oxygenated, deoxygenated, and total hemoglobin (HbO, HbR, HbT). **C**. Ratio maps averaged across 20 trials showing the response to a 2-s, 3-Hz train of air puffs to the right whisker pad. Maps show data from 1.7 s after stimulus onset except for the calcium map, which shows data from 0.1 s after stimulus onset. **D**. Average time course of the stimulus-evoked response in the contralateral barrel cortex (green region in panel C). The pink dashed line indicates when the calcium ratio map in panel C is shown whereas the black dashed line indicates when ACh, HbT and HbO ratio maps in panel C are shown. The grey-shaded area indicates the duration of the stimulus. **E**. Response of the contralateral barrel cortex to individual stimulus trains. The trials are sorted according to the magnitude of the calcium response.
